# Pharmacodynamic evaluation of AUM001/tinodasertib, an oral inhibitor of mitogen-activated protein kinase (MAPK)-interacting protein kinase 1, 2 (MNK1/2) in preclinical models and tissues from a Phase 1 clinical study

**DOI:** 10.1101/2024.02.23.581717

**Authors:** Bong Hwa Gan, Lay Hoon Lee, Reika Takeda, Maryam Yasin, Vincenzo Teneggi, Kantharaj Ethirajulu, Pauline Yeo, Dhananjay Umrani, Vishal Pendharkar, Darren Wan Teck Lim, Greg Li, Qingshu Lu, Yang Cao, Ranjani Nellore, Stephanie Blanchard, Hannes Hentze, Veronica Novotny-Diermayr

## Abstract

Mitogen-activated protein kinase (MAPK) interacting kinase (MNK) inhibitors affect cap-dependent mRNA translation by blocking the phosphorylation of RNA-binding proteins such as the eukaryotic initiation factor 4E (eIF4E). Phosphorylation on serine (Ser) 209 of eIF4E causes hyperactivation and dysregulation of mRNA translation, which is a hallmark of numerous malignancies. AUM001/Tinodasertib (ETC-206) is a selective and potent oral kinase inhibitor of MNK1 and MNK2 (IC_50_ of 64 and 86 nM, respectively), inducing dose-dependent inhibition of eIF4E phosphorylation on Ser209 (p-eIF4E) with an IC_50_ of 0.8 µM in K562-eIF4E cells. In mice, single oral doses of ∼12.5 mg/kg led to rapid (1-2 h post-dose) ∼70% inhibition of p-eIF4E in different normal or tumor tissues at a plasma concentration of 8.6 μM. However, in peripheral blood mononuclear cells (PBMCs), obtained from human healthy volunteers (HVs) in a Ph1 study, single oral doses of 10 or 20 mg ETC-206 did not show inhibitory activity up to 12 h post-dose, instead ETC-206 caused a statistically significant (p=0.0037) p-eIF4E inhibition in PBMCs of 24% at 24 h post-dose with 10 mg, and an inhibition of ≥27 % up to 52% was seen in 11/14 subjects in the 20 mg group where ETC-206 plasma concentrations remained above the IC_50_ for p-eIF4E (1.7 µM) for 30 h. While in mouse pharmacodynamic (PD) activity was also shown in tumor, skin, and hair follicles (HFs), in human tissues, PBMCs showed a trend for delayed PD inhibition and skin was not a suitable surrogate. Analysis of pharmacokinetic (PK) and PD relationships shown herein demonstrate excellent pharmaceutical properties of ETC-206 which has now advanced to Ph2 clinical trials (NCT05462236).

## Introduction

MNK1 and MNK2 are serine-threonine kinases of the MAPKs family [1] which are activated via phosphorylation on threonine (Thr) 197/202 by the MAPKs ERK and p38. MNKs are the sole kinases known to phosphorylate the eIF4E on Ser209 [2], and are a part of the large eIF4F complex, playing a central function in cap-dependent mRNA translation. MNKs need to be in physical proximity to eIF4E for the Ser209 phosphorylation to take place, which is achieved by binding to the scaffolding protein eIF4G, another component of the eIF4F complex. The affinity of MNKs to eIF4G in turn is regulated by a Pak2-mediated phosphorylation of MNK1 on Thr22/Ser27 [3], thereby adding an additional level of regulation to the phosphorylation of eIF4E [4, 5].

MNK inhibitors were initially of interest because eIF4E was described as a bona-fide oncogene with phospho-(p)-Ser(S)209-eIF4E being necessary for tumorigenesis but dispensable for normal development [6–9]. Knock-in studies with a S209A mutant of eIF4E further confirmed that a loss of Ser209 phosphorylation does not cause developmental defects or a loss of cell viability [10], while a multitude of independent studies demonstrated that increased levels of p-S209-eIF4E were linked to poor prognosis in a variety of different cancers [11, 12], promoted epithelial to mesenchymal transition, invasion and metastasis [13]. However, the interest in targeting MNKs has picked up because it was noted that a decrease of p-S209-eIF4E due to MNK inhibition specifically and disproportionally affecting the translation of a subset of mRNAs that promote cell survival, cytokines, tumor growth and invasion, while not affecting global protein translation. Among the differentially translated proteins in tumors is the programmed death ligand 1 (PD-L1) [9, 10, 14, 15]. Recently, studies demonstrated that cap-dependent mRNA translation plays an important role in a multitude of cells of the tumor microenvironment with levels of p-S209-eIF4E causing differential effects in neutrophils, macrophages, regulatory T-cells and Th1 cells and T-cells, affecting cell survival and cytokine production [16–18], giving rise to the hypothesis that MNK inhibitors in combination with immunotherapy will potentially provide effective treatment options for cancer. Tomivosertib (eFT508; Effector Therapeutics Inc.) was the first highly selective small molecule inhibitor of MNK1/2 with *in vitro* potency in the nanomolar range that entered into clinical studies [19]. In preclinical mouse models, it was demonstrated that liver tumors with a particular molecular phenotype showed a 50% reduction in p-eIF4E and PD-L1 levels after treatment with 10 mg/kg eFT508 [15]. In a Ph2a study it was demonstrated that the combination of eFT508 with PD-1/PD-L1 inhibitors led to a significant increase in progression-free survival in non-small cell lung cancer (NSCLC) patients. Recently a randomized, double-blinded, placebo controlled pivotal study has been started in the same indication [20].

ETC-206 is a small molecule MNK1/2 inhibitor of that is highly selective for MNK kinases, with an *in vitro* IC_50_ of 64 and 86 nM for MNK1 and MNK2, respectively. ETC-206 exhibits favorable PK parameters and induced tumor regression in combination with BCR-Abl inhibitor (dasatinib) in a murine model of blast crisis chronic myeloid leukemia (CML) [1]. ETC-206 was initially assessed in a single ascending dose trial, followed by a multiple ascending dose trial, performed in human HVs. ETC-206 was shown to be safe and well tolerated [21], without any treatment-emergent adverse events or dose limiting toxicities and is now being explored in an ongoing Ph2 trial in metastatic colorectal cancer in combination with either pembrolizumab or irinotecan (NCT05462236). Here, we describe the use of a Western blot assay to demonstrate target engagement, assess PK/PD relationships in preclinical models and compare to results obtained in a first-in-human (FIH) HV study after ETC-206 single dose.

## Material and Methods

### Compounds

ETC-206 (Syngene International Ltd, Bengaluru, India) was prepared for *in vitro* studies as a 10 mM stock in DMSO. For oral administration in animals, ETC-206 was prepared in sodium carboxymethyl cellulose (Na-CMC) containing 0.5% Tween^®^ 80, to yield a 99.5% (w/v) Na-CMC suspension that was vortexed, sonicated, and dosed at 10 mL/kg. ETC-206 for human oral ingestion was formulated as 10 mg strength in white, opaque capsules with matching placebo capsules and manufactured at Patheon UK Ltd (Milton Park, United Kingdom).

### Phase 1 healthy volunteer study

Biomarker samples (HFs, PBMCs, skin biopsies) were collected from a total of 23/24 subjects recruited in a Ph1 study conducted in Singapore (IRB approval #2016/2522) described earlier [21]. Participants provided informed consent before starting any study-related procedure. See *Supplementary Methods* for details of sample collection/processing.

### Cell lines, primary cells, and cell lysis

The human CML cell line K562 (#CCL-243^™^, American Tissue Type Collection [ATCC]) was batch-transfected with the MSCV-IRES-GFP vector ENREF76 [9], encoding full-length murine eIF4E (named K562-eIF4E). Transfected cells were FACS-sorted for cells expressing high levels of eIF4E. Cells were grown according to ATCC’s instructions and authenticated by Axil Scientific (Singapore). PBMCs were obtained from healthy donors under IRB approval #NUS2110 or during the ETC-206 clinical study (IRB approval #2016/2522), collected using Vacutainer CPT^™^-tubes (#362761, Becton, Dickinson and Company), washed and plated in 6-cm cell culture dishes in RPMI 1640 containing 10% FBS and 1% penicillin-streptomycin solution, and maintained at 37°C and 5% CO_2_ for 2-3 h or overnight prior to treatment. Cells were lyzed in Complete Lysis Buffer on ice, prepared by adding HALT^™^ protease and phosphatase inhibitor cocktail (#78440) to a final of 1X concentration to Pierce^®^ RIPA buffer (#89900), from Thermo Fisher Scientific.

### Animal studies

All rodents (InVivos, Singapore) were housed in individual ventilated cages under controlled conditions, in compliance with NIH and NACLAR guidelines. Experiments were performed under IACUC Approval #151001. Mice between 6 and 12 weeks of age were used. PK/PD studies were performed in non-tumor-bearing female ICR mice (IcrTac:ICR or Hsd:ICR(CD-1^®^), or in female SCID mice (C.B-IgH-1^b^/IcrTac-Prkdc^scid^) bearing K562-eIF4E xenografts as described previously [1]. After ∼12 days, when tumors reached a volume of ∼200-500 mm^3^, animals received a single dose of ETC-206, and tissues were harvested at different time points post-dose. Tumor volumes were calculated in mm^3^ using w^2^×l/2 (w = width, l = length in mm).

### Tissue homogenization

Briefly, tissues (tumor, HFs, skin) were homogenized in cold Complete Lysis Buffer containing either zirconia or stainless steel beads (Tomy Digital Biology Co Ltd.) using Micro Smash^™^ tissue homogenizer (Tomy Digital Biology Co Ltd.) at a speed of 4,000 rpm under external cooling using dry ice in 5 cycles of 10 s each (tumor, skin) or 3 cycles of 5 min each (HFs) followed by 1 min of cooling between cycles. After 5 min incubation on ice, lysates were cleared by a 15 min spin. Human tissues were collected from HVs in the Ph1 study previously described [21]. For detailed methods used to prepare lysates from mouse and human samples see *Supplementary Methods*.

### Western blot

Total protein concentrations were determined against a standard curve of BSA in a 96-well plate using the DC^™^ Protein assay (Bio-Rad Laboratories). Proteins were separated on reducing 10% NuPAGE^™^ Novex^™^ Bis-Tris Protein Gels and transferred onto PVDF membranes, blocked (5% BSA in 1X TBST containing 0.05% Tween-20) at room temperature, and probed overnight at 4°C with either of the following antibodies: p-S209-eIF4E (#9741), total eIF4E (#2118), all diluted 1:1,000; or GAPDH clone 14C10 (#2118) diluted 1:10,000. Secondary antibodies used were anti-rabbit-IgG-HRP or anti-rabbit-IgG-AP, (#7074, #7054) diluted 1:2,000. All antibodies were purchased from Cell Signaling Technology. For developing, either Amersham ECL^™^ Prime Western Blotting Detection Reagent (Cytiva) or Immune-Star^™^ AP Substrate (Bio-Rad Laboratories) were used. Images were digitally captured using Amersham Imager 600 UV (Cytiva) and analyzed densitometrically using the ImageQuant^™^ TL software v8.1 (Cytiva). Relative p-eIF4E levels were determined as a ratio of p-eIF4E levels over total eIF4E measured in the same sample. To determine the percentage inhibition relative p-eIF4E levels in a sample were compared to relative levels of p-EIF4E from the same subject pre-dose (or vehicle-treated mice), set to 100%.

### Bioanalysis and PK

Plasma samples were collected using K_2_EDTA as anti-coagulant. Dosing solutions of preclinical experiments were diluted 8,000-fold in a 1:1 mixture of methanol and water. Mouse tissue samples (plasma, skin, tumor) were diluted 10-fold. Detailed methods for extraction and bioanalysis are provided in the *Supplementary Methods.* Bioanalysis of human plasma samples was performed blinded at SGS (Cephac, France), using an analytical validated method under GLP. PK parameters of human samples were analyzed by Phinc Development (Évry, France) using Phoenix WinNonlin^®^ Professional software v6.4, Pharsight Corporation).

### Statistical analyses

Statistical analyses (if not otherwise mentioned) have been performed using GraphPad Prism 7 for Windows, v7.02 (GraphPad Software Inc.). For clinical sample analysis normality was tested using the D’Agostino and Pearson omnibus normality test. Sphericity was not assumed. For PBMCs repeated measures one-way ANOVA was performed using all time points that did not have missing data, to determine significance of inhibition and to test for linear trend of inhibition. Where the data was not normally distributed, significance was determined using the Wilcoxon Matched Pairs Signed Rank Test or the values were transformed (log_2_) prior to performing the analysis.

## Results

### Determination of the IC_50_ for inhibition of p-eIF4E by ETC-206 in human cells *in vitro*

ETC-206 is a potent and selective MNK1/2 inhibitor, which was confirmed in a SelectScreen**^®^** panel with 414 recombinant human kinases, where only 38 kinases were identified that were inhibited >50% at 10 µM. IC_50_ determination using the same assays for each of these 38 kinases confirmed that ETC-206 is most active against MNK1 (IC_50_: 64 nM) and MNK2 (IC_50_: 86 nM), with the next lowest IC_50_ of 610 nM (S1 Table) for receptor interacting serine/threonine kinase 2 (RIPK2). ETC-206 was mildly anti-proliferative when tested using CellTiterGlo^®^ luminescent cell-viability assay in a panel of 71 tumor and non-cancerous human cell lines, with cellular IC_50_ mostly in the 10-45 µM range, and only 5 cell lines showed inhibition <10 µM (S2 Table).

To establish the p-eIF4E inhibition as a PD target engagement biomarker, K562-eIF4E cell line and HVs’ PBMCs were treated *in vitro* for 2 h with 12 nM to 50 μM ETC-206, and p-eIF4E, eIF4E and GAPDH levels were determined by Western blot. Treatment with ETC-206 led to a concentration-dependent inhibition of relative p-eIF4E levels, where 50 μM ETC-206 led to a ≥90% inhibition (Fig 1). The IC_50_ of p-eIF4E was determined to be 0.8 μM for K562-eIF4E cells and about 2-fold higher (1.7 μM) in primary human PBMCs. Concentrations below ∼0.8 μM (K562-eIF4E cells) or ∼0.5 μM (PBMCs) only gave a marginal p-eIF4E inhibition. Normalizing p-eIF4E levels using total eIF4E levels gave comparable results to normalization using GAPDH levels, the latter was therefore not determined for all following studies.

**Fig 1.**
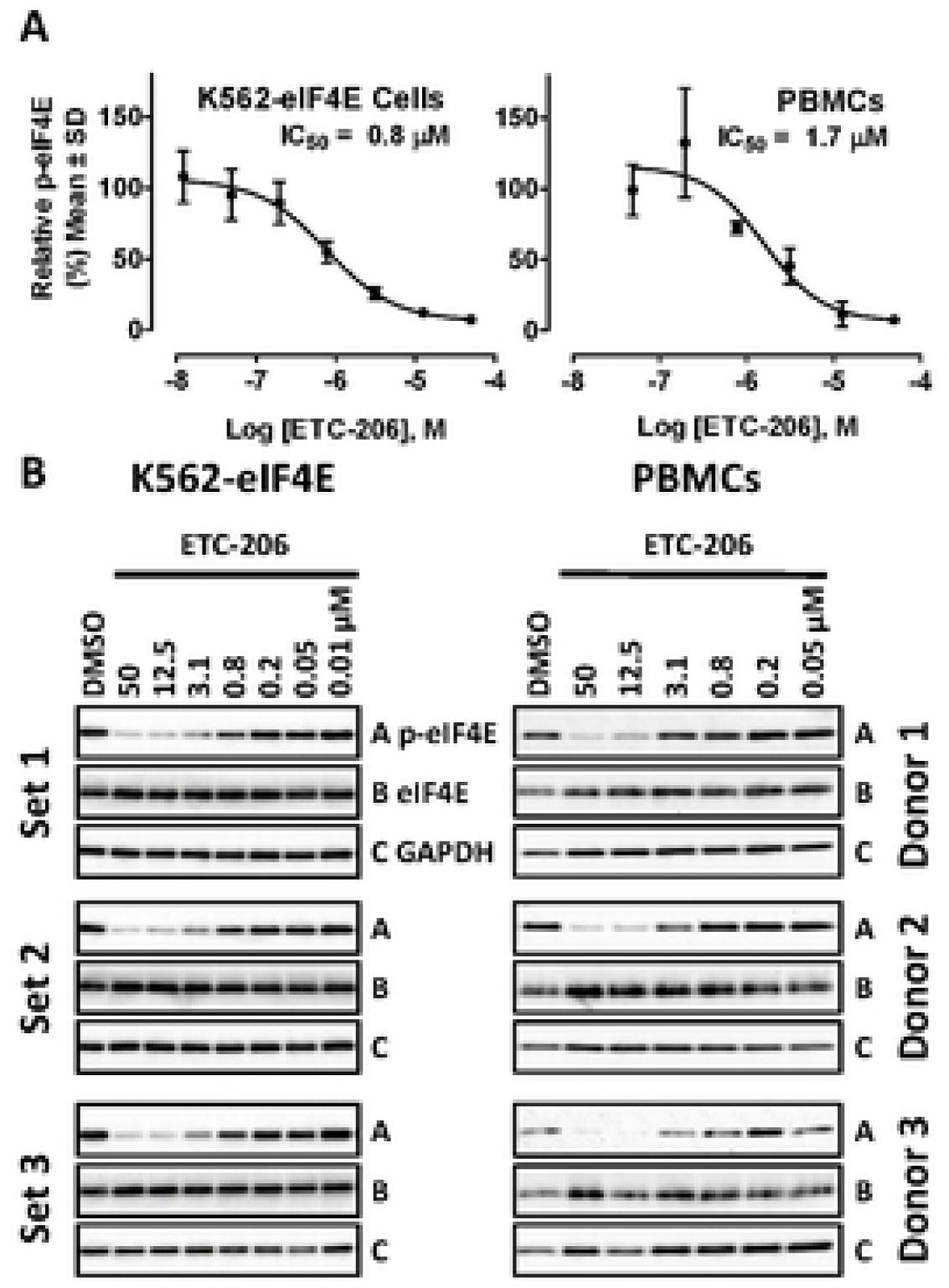
ETC-206 inhibits p-eIF4E in human cells *in vitro*. K562-eIF4E cells or primary human PBMCs obtained from volunteer donors were treated with different concentrations of ETC-206 or vehicle control (0.1% DMSO) as indicated for 2 h. Cells were lyzed, separated on Bis-Tris protein gels, and Western blot analysis was performed using antibodies against p-eIF4E, eIF4E, and GAPDH as indicated. **(A)** IC_50_ curves showing relative p-eIF4E levels in K562-eIF4E cells and in PBMCs. Relative p-eIF4E levels were determined densitometrically Western blots to calculate IC_50_ values; For K562-eIF4E cells n=6; “Set 1” refers to one sample set from those 6 replicates. **(B)** Corresponding Western blots from one half of the experiments (n=3) is shown; 5 μg/10 μg of protein loaded per lane for K562-eIF4E cell line/PBMCs, respectively.

### Effects of ETC-206 on inhibition of p-eIF4E in mouse normal tissues

The inhibitory effect of ETC-206 on relative p-eIF4E levels in different normal (non-cancer) tissues was determined in ICR mice (Fig 2). Comparing tissue samples from mice treated for 1 h with a single dose of 200 mg/kg ETC-206 to vehicle-treated mice, an inhibition of p-eIF4E of 53% in bone marrow, 62% in PBMCs, 82% for skin, and 78% in HFs was determined (Fig 2A). In platelets, no p-eIF4E inhibition could be determined as expression of eIF4E was very low compared to other tissues (Fig 2A, bottom panel). PBMCs, skin, and HFs were chosen for further analysis of p-eIF4E inhibition.

**Fig 2.**
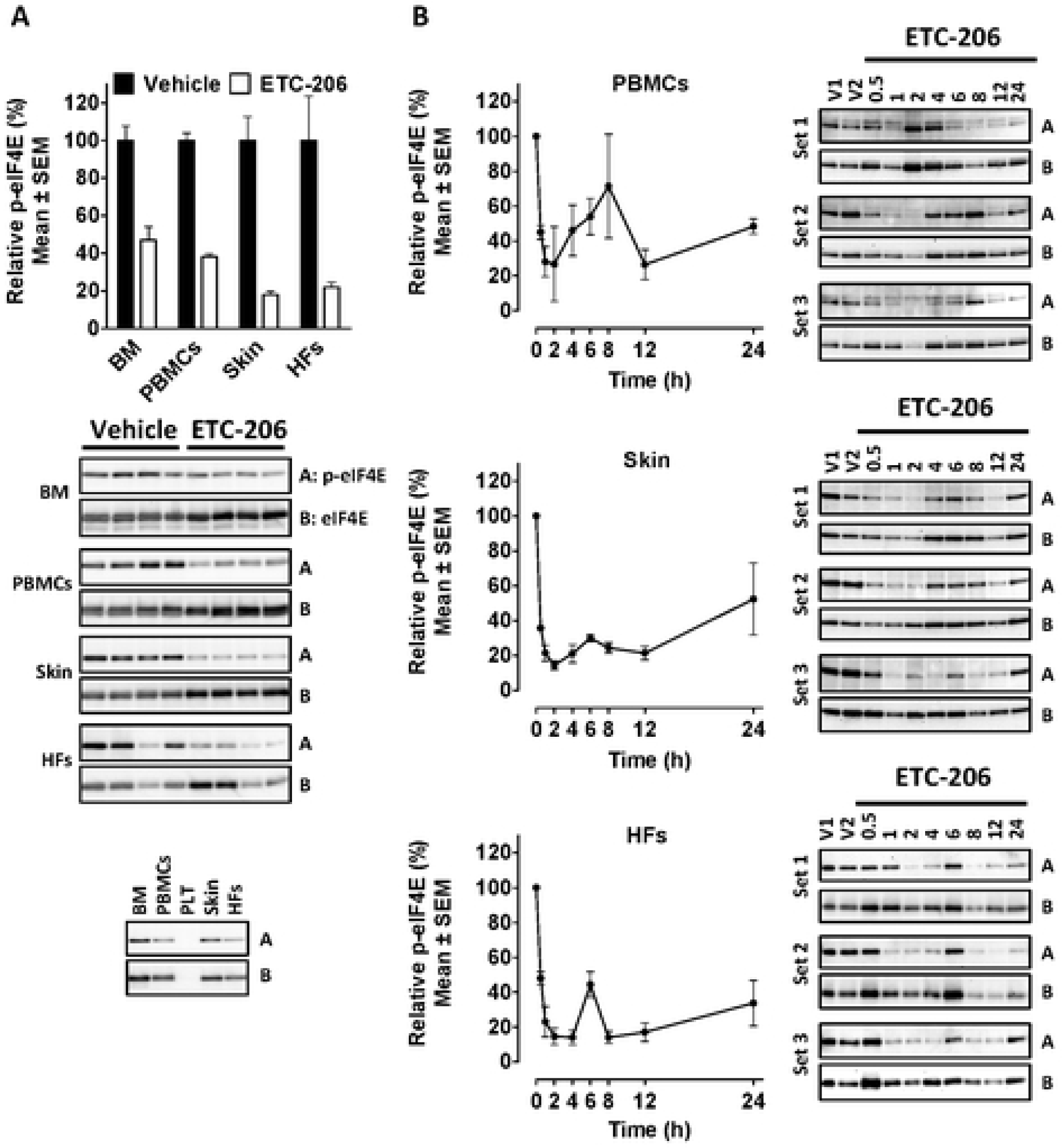
Inhibition of eIF4E phosphorylation by single-doses of ETC-206 in surrogate tissues from ICR mice. Female ICR mice were dosed orally with a single dose of vehicle or 200 mg/kg ETC-206 (n=4/group). (A) Mice were euthanized, and tissues collected 1 h post-dose. Tissue samples from surrogate tissues were analyzed via Western blot analysis with antibodies as indicated (BM: bone marrow, 15 μg protein per lane; PBMCs: peripheral blood mononuclear cells, 30 μg protein per lane; skin, 15 μg protein per lane; HFs: hair follicles, 12 μg protein per lane). **(A)** Top panel: densitometric analysis of relative p-eIF4E levels; Middle panel shows corresponding Western blots; Lower panel: equal amounts (12 μg protein) of tissues as indicated (PLT: platelets) were loaded to compare protein levels in different surrogate tissues. **(B)** Mean relative p-eIF4E levels in surrogate tissues from ICR mice collected at different time points post-dose (0.5 to 24 h) were assessed by Western blot; corresponding blots are shown on the right (V1: pooled vehicle samples (n=3) used to calculate 100%, V2: single sample from one vehicle-treated animal; “Set 1” refers to one sample set from those 3 replicates. (5 μg protein per lane for PBMCs, skin; 3 μg for HFs).

The time point of maximum inhibition and the duration of p-eIF4E inhibition in ICR mice were determined after receiving a single dose of 200 mg/kg ETC-206. Relative p-eIF4E levels were measured at different time points post-dose in PBMCs, skin, and HFs (0.5-24 h), and compared to relative p-eIF4E levels of vehicle-treated mice, considered to be 100% at 0 h (Fig 2B). The onset of the inhibitory effect was rapid with 55-65% inhibition observed in all tissues at 0.5 h post-dose and maximum inhibition achieved at 1-2 h post-dose (∼85% in skin and HFs, ∼70% in PBMCs). High levels of inhibition were sustained for up to 12 h, and 24 h after a single-dose treatment with ETC-206, p-eIF4E inhibition was still between 50-65% in all three tissues when the plasma concentration was around ∼5,000 ng/mL or 12 µM (data not shown) in an independent experiment. Overall, a higher variability of p-eIF4E inhibition was observed in PBMCs compared to other tissues tested.

### Effective dose range to inhibit p-eIF4E in different tissues *versus* plasma exposure after a single dose of ETC-206 in mice

To determine the minimal inhibitory dose of ETC-206 and the corresponding plasma concentrations *in vivo*, ICR and SCID mice received single oral doses of ETC-206, and the relative p-eIF4E levels in PBMCs, skin, HFs, and tumor samples (for SCID mice carrying K562-eIF4E tumors) were compared to vehicle-treated mice 2 h post-dose, with ETC-206 plasma concentrations analyzed in the same animals. ICR mice received 8 doses of ETC-206 (1-200 mg/kg) and SCID mice received 5 doses (12.5-200 mg/kg). ETC-206 plasma levels showed an overall similar linear dose-exposure relationship in ICR and SCID mice across the dose range tested with slightly higher ETC-206 exposure in ICR mice (Fig 3A). In ICR mice, a rapid dose-dependent decrease in relative p-eIF4E levels was observed, with a minimum inhibitory dose of approximately 12.5 mg/kg in skin, HFs and spleen (Fig 3B upper panels and S1C Fig). In PBMCs, variation was high and only the highest dose of 200 mg/kg resulted in significant p-eIF4E inhibition of ∼70% (Fig 3B). In skin and PBMCs, lower doses of ETC-206 led to an increase in p-eIF4E levels beyond basal levels (∼1.7-fold at the doses of 1-3 mg/kg for skin, ∼2.5-fold at 6-25 mg/kg in PBMCs), yet, there was a statistically significant, strongly negative correlation between ETC-206 plasma concentrations and p-eIF4E levels in all tissues except PBMCs (Pearson’s r between -0.75 and -0.77 compared to -0.38 in PBMCs; Fig 3B, lower panels).

**Fig 3.**
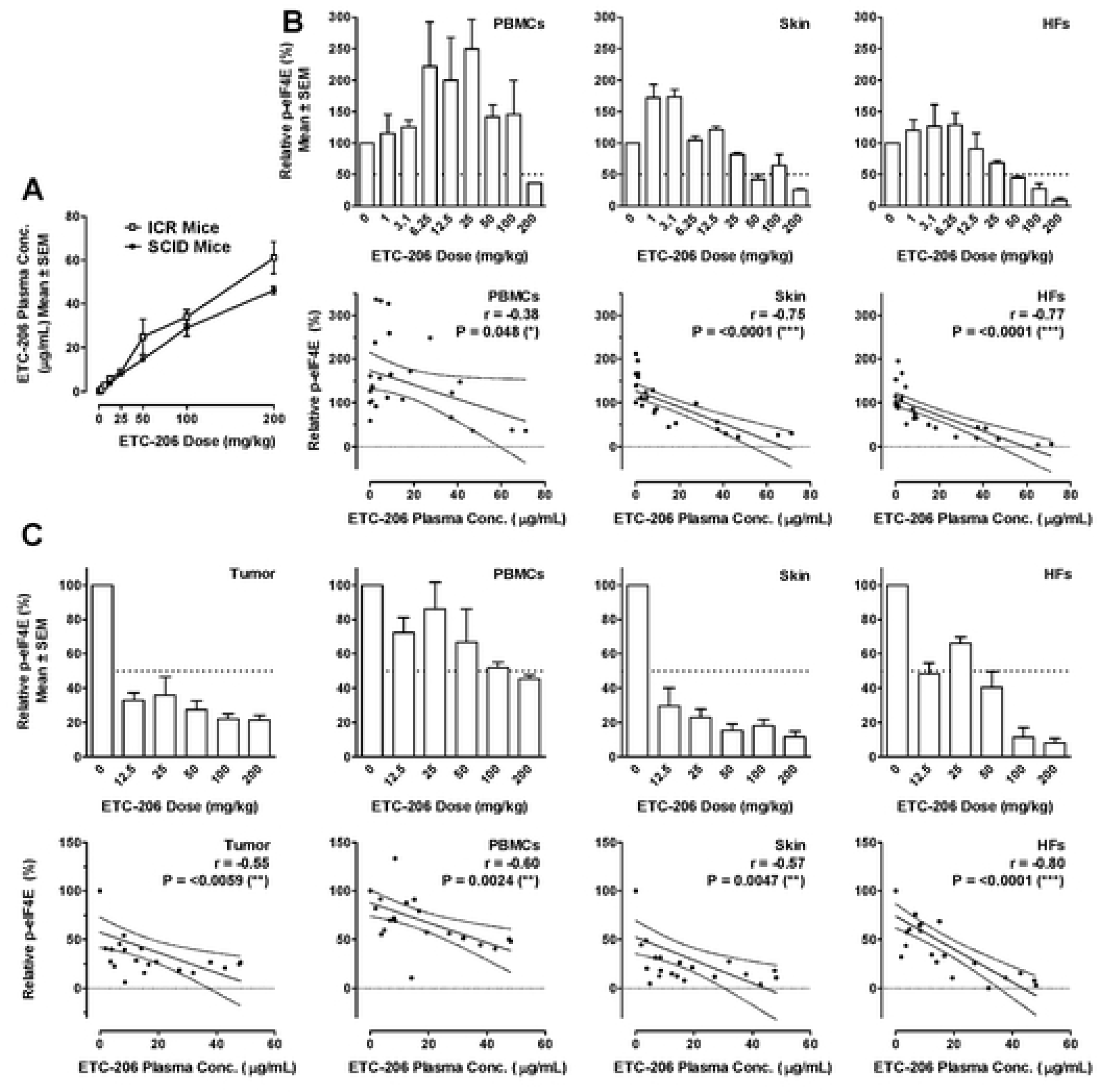
Plasma concentrations, relative p-eIF4E levels, and PK/PD correlations in tissues from ICR mice and tumor-bearing SCID mice after a single-dose of ETC-206. **(A)** Plasma concentrations of ETC-206 were determined by UPLC-MS/MS 2 h post-dose in female ICR (n=3/dose) and tumor-bearing female SCID mice (n=4/dose). Dose levels tested were 1, 3.1, 6.25, 12.5, 25, 50, 100, 200 mg/kg for ICR mice, 12.5, 25, 50, 100, 200 mg/kg for SCID mice. **(B)** Upper panels: mean relative p-eIF4E levels from surrogate tissues of ICR mice treated with ETC-206 for 2 h at the indicated dose levels were determined by Western blot followed by densitometry. Lower panels; correlation between ETC-206 plasma concentrations and p-eIF4E inhibition in individual animals (n=27); solid line: linear correlation, dotted lines: 95% confidence intervals, Pearson’s r and corresponding P values are shown; **(C)** Upper panels and lower panels as described under (A) but in SCID mice bearing K562-eIF4E tumor (n=4/dose). Tumors collected had a mean volume of 422 mm^3^ 26 days post implantation.

Next, we assessed if p-eIF4E levels in tumor could be inhibited at a lower dose compared to non-tumor tissue. Subcutaneous K562-eIF4E tumors were grown in SCID mice as previously described [1]. In this mouse model of human blast crisis chronic myeloid leukemia (CML) ETC-206 was previously shown to have anti-tumor activity when dosed in combination with a sub-optimal dose of dasatinib [1]. Tumors and normal tissues (HFs, skin, PBMCs) were harvested after single doses of 12.5-200 mg/kg ETC-206. In tumor-bearing SCID mice, relative p-eIF4E levels at 2 h post-dose were significantly inhibited in all tissues (tumor, PBMCs, HFs, skin) with a maximum inhibition of ∼90% at the highest dose tested (200 mg). Tumor and skin were the most sensitive tissues, reaching ∼70-75% inhibition at the lowest dose tested (12.5 mg/kg). Inhibition of p-eIF4E was significantly negative correlated with plasma concentrations for all tissues tested (-0.55≤r≤-0.80, Fig 3C). However, p-eIF4E inhibition in tumor tissue was similar to that seen in skin at 12.5 mg/kg. PBMCs again showed highest variability in relative p-eIF4E levels.

In summary, ETC-206 showed a linear dose-exposure relationship in both mouse strains tested, and a significant (≥70%) inhibition of p-S209-eIF4E was seen in SCID mice with 12.5 mg/kg (corresponding to ≤3,500 ng/mL or 8.6 μM). The observed inhibition in any tissue was never complete but on average around 80% from baseline levels. Maximal inhibition in all preclinical tissues was seen between 2-4 h post-dose.

### Inhibition of p-eIF4E in human tissues

The same assay established using preclinical tissues was used to evaluate PD in clinical samples collected in the single ascending dose Ph1 FIH study in male, human HVs, as previously described [21]. ETC-206 was dosed in three different dosing periods (DPs) at 10 mg or 20 mg in the fasted state (DP1 or DP2, respectively), or at 10 mg in a fed state (DP3) to a total of 23 HVs as described [21]. PBMCs and HFs were collected at all DPs at up to 9 different time points (0-30 h). In DP3 additional pre-dose and a 1.5 h post-dose skin samples were analyzed from 9 donors.

In contrast to mice, no inhibition of p-eIF4E was observed in human PBMCs within the first 4 h after a single dose, however, inhibition was seen from 24 h onwards. At DP1, relative p-eIF4E levels showed an inhibition of 24%, which was statistically significant (P=0.0037; Fig 4A; S2A Fig). The heat map shows that at 24 h post-dose 11/16 subjects treated showed ≥15% and up to 56% inhibition (Fig 4B). There was a linear trend for inhibition with a slope of -0.036 that was significant (p=0.0043). At DP2, an inhibition of ≥27% up to 52% post-dose was seen with 8/14 subjects at 24 h, and ≥12% up to 70% in 11/14 subjects at 30 h post-dose. There was a statistically significant linear trend for inhibition with a slope of -0.051 (p<0.0001). Similarly, for DP3, the maximum inhibition was seen at 30 h post-dose with inhibition ≥24% up to 71% in samples from 5/9 subjects at 30 h but without linear trend for inhibition. In PBMCs of placebo treated subjects no mean inhibition was observed at any time point.

**Fig 4.**
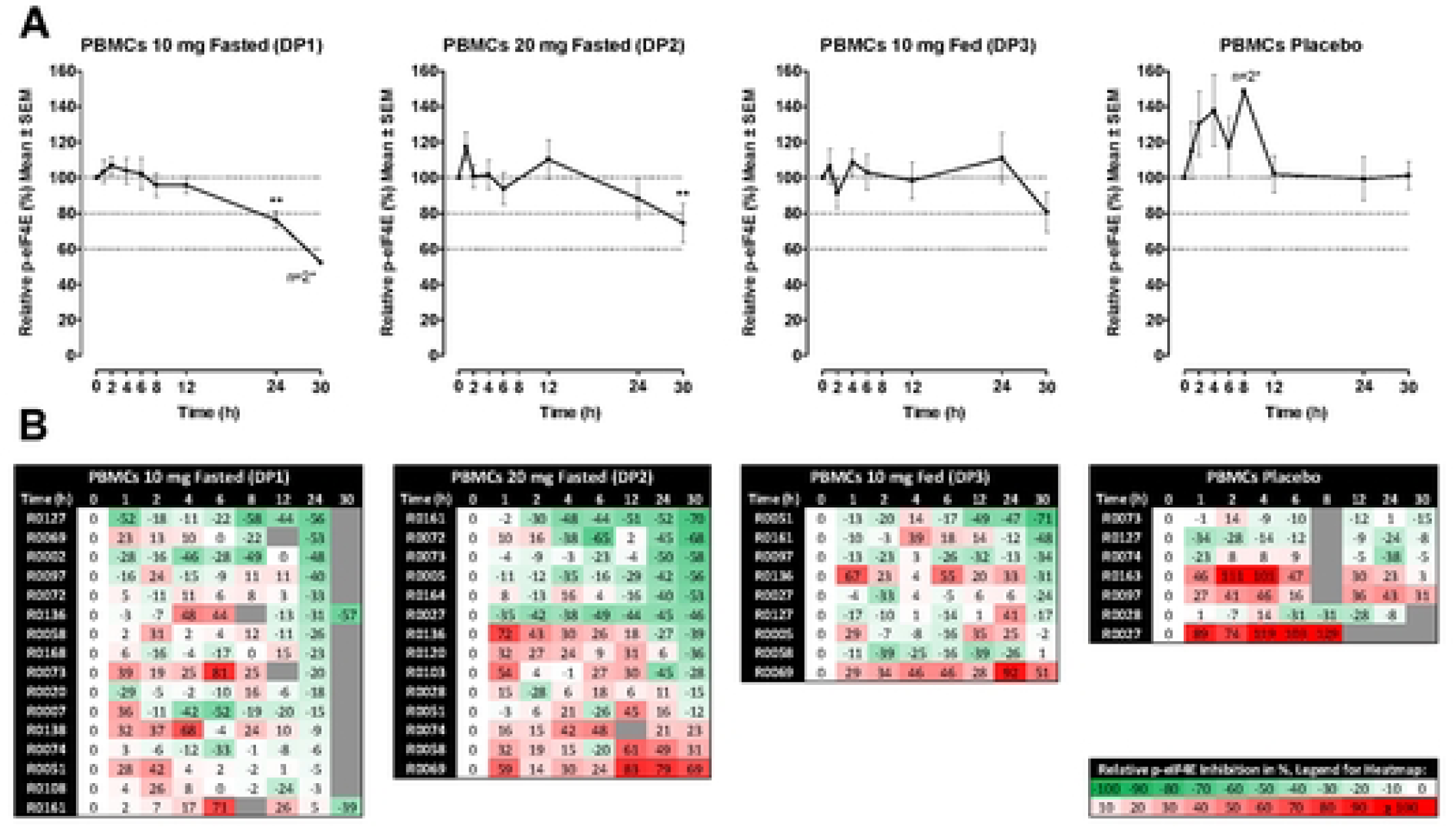
Effects of a single-dose of ETC-206 on relative p-eIF4E levels in PBMCs from human volunteers. Subjects (male HVs) were dosed with 10 mg (n=16) or 20 mg (n=14) ETC-206 in a fasted state, or with 10 mg (n=9) ETC-206 in fed state or received placebo (n=7). Whole blood was collected in CPT-vacutainers tubes with sodium citrate as anti-coagulant, at time points as indicated from pre-dose (0) to 24 h or 30 h (± 6 min). PBMCs were isolated and immediately snap-frozen. Relative p-eIF4E levels were determined in PBMCs as described for Fig 3B. **(A)** Mean relative p-eIF4E levels are shown per time point, for a time point with only n=2 no SEM is shown (as indicated by *). ** indicates P <0.005, for the one-way ANOVA with Geisser-Greenhouse correction. **(B)** Heat map of relative p-eIF4E levels in PBMCs per dosing group and shaded according to the inhibition, see inserted legend; Grey fields without numbers indicate samples not collected, samples lost during analysis, or outliers.

In HFs at DP2, there was an approximately 20% inhibition observed from 6 h onwards, but no significant inhibition seen in HFs from subjects treated with 10 mg or placebo (Fig 5A and 5B; S2B Fig). Human skin, in contrast to murine skin, had much lower levels of total eIF4E and no change in the low amounts of p-eIF4E levels was observed. In addition, the observed protein size for p-eIF4E in human skin was variable (S2C Fig).

**Fig 5.**
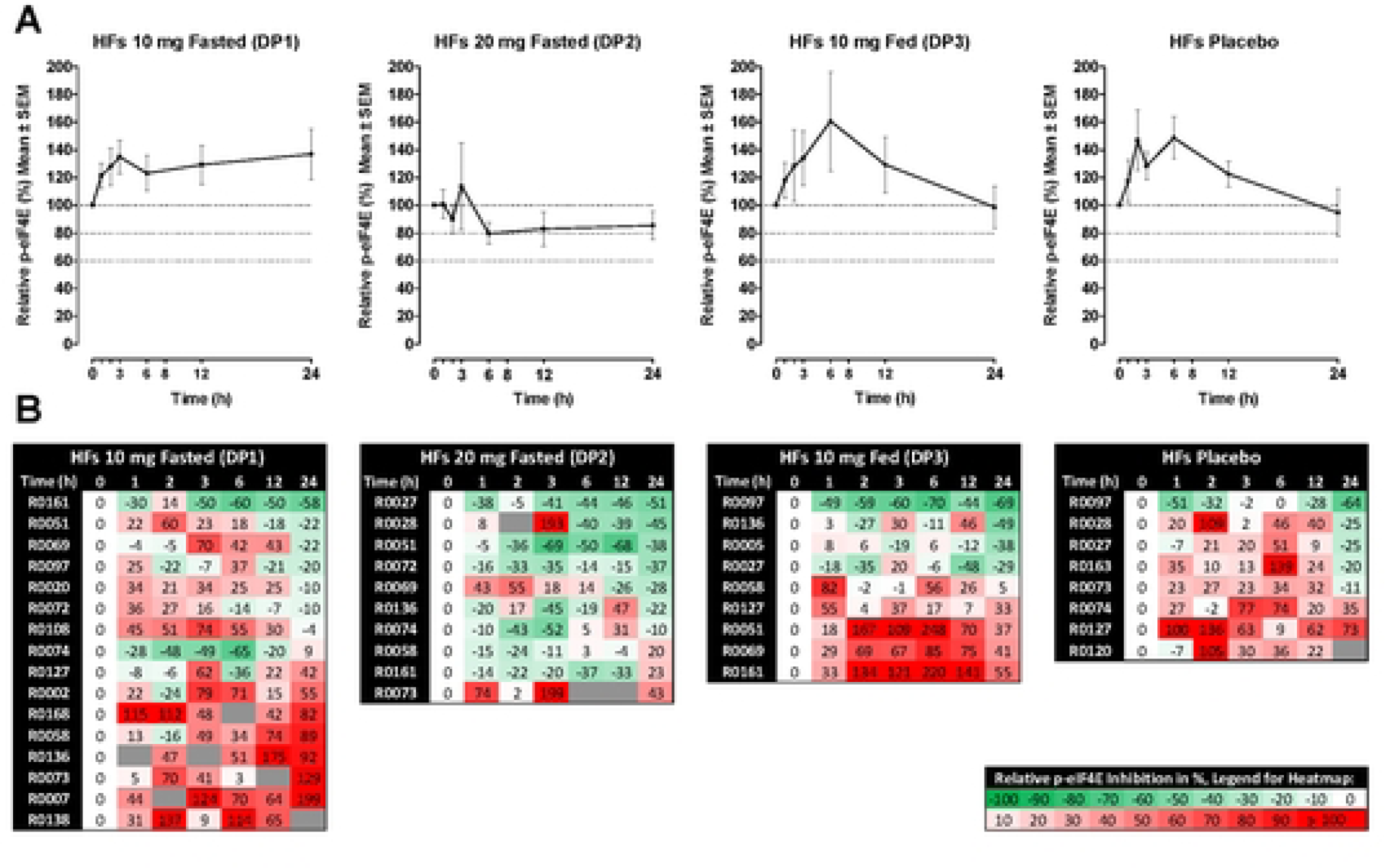
Effects of a single-dose of ETC-206 on the inhibition of relative p-eIF4E levels in hair follicles from human HVs. At least 40 hair follicles (HFs) were collected at the indicated times (±15 min). Western blot and densitometry were performed as described for Fig 3B, Samples from 4 volunteers had to be excluded from all analyses as the pre-dose sample did not yield any protein (not shown). For **(A)**, **(B)**, see legend for Fig, 4.

### Comparing murine and human PK/PD relationships of ETC-206

Fig 6A shows the human plasma PK of ETC-206 over 72 h (for DP1) and over 144 h for the other two dosing groups (DP2, DP3), with sampling time points for biomarkers indicated by vertical lines. The T_max_ for the two groups dosed in fasted state (DP1, DP2) was about 1 h post-dose. The mean C_max_ for the 10 mg groups (DP1, DP3) was close to 1 µg/mL, and reached about 2 µg/mL for the 20 mg group (DP2). A weak, non-significant negative correlation between C_max_ and p-eIF4E inhibition at 24 h post-dose was found when all individual data pairs were analyzed (DP1-DP3 and placebo, n=46), similarly the correlation between AUC_0-∞_ and p-eIF4E inhibition was slightly negative, but not statistically significant (Fig 6B).

**Fig 6.**
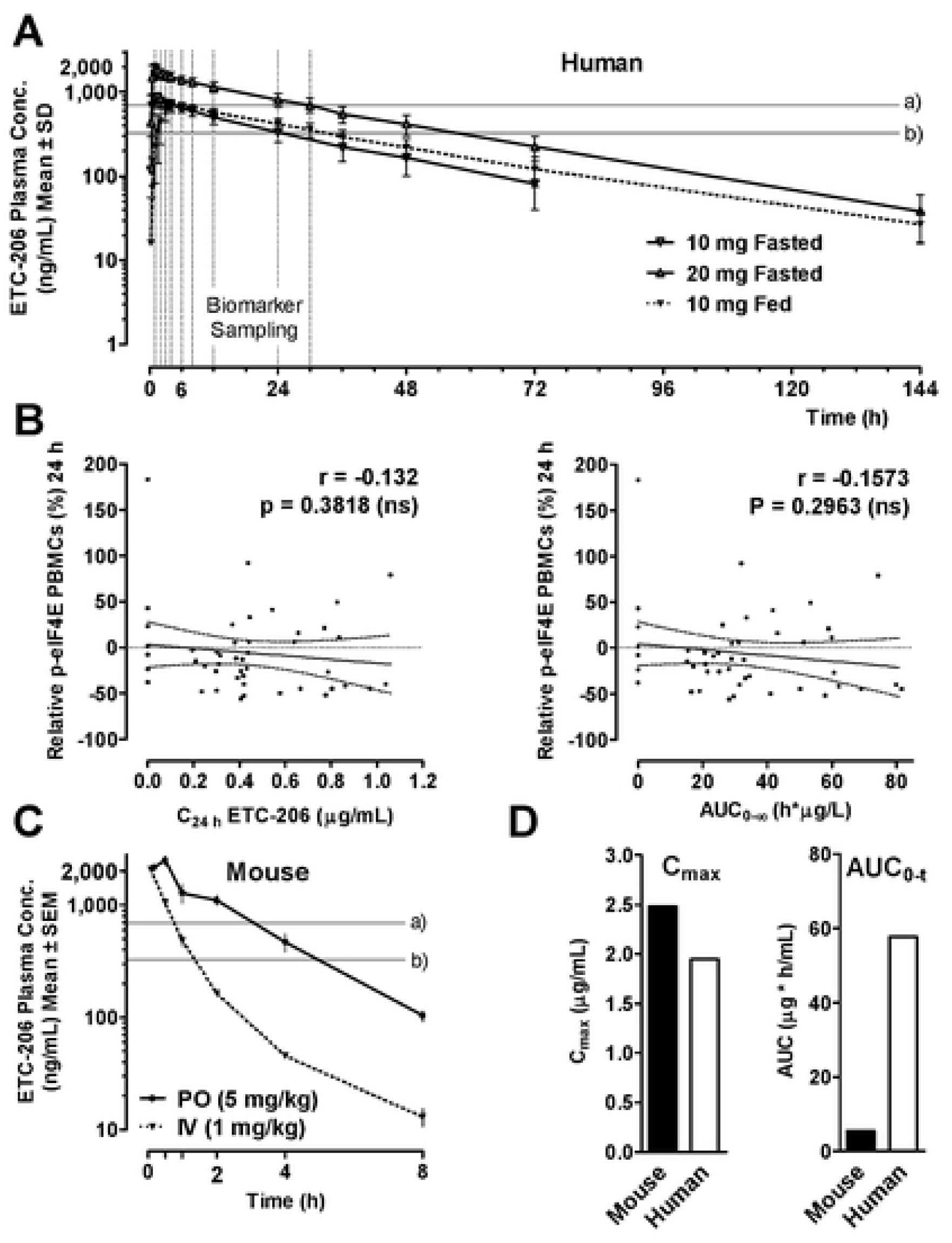
PK/PD comparison of ETC-206 in human HVs and ICR mice. (**A**) PK of ETC-206 in human plasma, indicating the time points of biomarker sampling (dotted vertical lines) For 10 mg fasted (DP-1): n=16, except for the 30 h and the 144 h time point (n=2); for 20 mg fasted (DP2): n=14, except for 144 h (n=11); and for 10 mg fed group (DP3)] n=9, except for 144 h (n=7). The IC_50_ values for inhibition of eIF4E phosphorylation in human PBMCs (694 ng/mL), line a, and in K562-eIF4E cells (327 ng/mL), line b, are shown. **(B)** Correlation between relative levels p-eIF4E inhibition and plasma concentration at 24 h post-dose (left) or AUC_0-infinity_ (right) in PBMCs at 24 h (n=46, lines, r and P: as described in Figure 3). **(C)** ETC-206 plasma concentrations were determined in ICR mice after oral (PO) or intravenous (IV) dosing of fasted at the time points indicated (n=3/group). Lines a) and b) show the same p-eIF4E IC_50_ values as in Figure A. **(D)** Comparison of C_max_ and AUC_0-t_ in humans dosed with 20 mg ETC-206, fasted state) *versus* ICR mice, dosed at 5 mg/kg PO, ETC-206 (non-fasted).

The PK profile of ETC-206 was compared between mouse and human using doses that are roughly comparable (5 mg/kg dose in mice is the human equivalent dose of 20 mg, Fig 6A, 6C). The C_max_ levels observed in human were below the 3,500 ng/mL threshold shown to cause acute p-eIF4E inhibition in SCID mice, but the plasma concentration remained above the *in vitro* IC_50_ for p-eIF4E inhibition in PBMCs of 1.7 µM (694 ng/mL) for 30 h (see Fig 6A). Due to a 14-fold higher mean half-life in human (Fig 6A, t_1/2_ = 25.8 h) compared to mice (Fig 6C, t_1/2_ = 1.8 h), the mean AUC_0-t_ was about 10x greater in human at comparable C_max_ (Fig 6D), which may explain the delayed p-eIF4E inhibition observed in the human study.

## Discussion

Blocking cap-dependent mRNA translation selective and potent inhibitors of MNK has proven to provide clinical benefits for cancer patients with eFT508 being granted orphan drug status by FDA for further development in diffuse large B-cell lymphoma [19] and having shown favorable clinical results in the Ph2 trial in NSCLC in combination with pembrolizumab [20]. There is strong evidence that MNK inhibition blocks multiple oncogenic proliferation pathways, reduces pro-inflammatory and pro-tumorigenic cytokines, while increasing anti-tumor immunity [22–24]. While cytotoxic activity has been described for non-specific MNK inhibitors *e.g.*, cercosporamide or CGP57380, selective MNK inhibitors *e.g.*, ETC-206, eFT508 and BAY 1143269 have very little anti-proliferative activity with cellular IC_50_ >30 µM in a majority of cell lines (S1 Table) [24, 25].

Besides ETC-206, only eFT508 and BAY 1143269 entered the clinic with BAY 1143269 being similar to ETC-206 in terms of potency for MNK and selectivity (MNK1 IC_50_ 40 nM, MNK2 IC_50_ 904 nM and PIM1 IC_50_ 518 nM), while eFT508 is a more potent MNK1/2 inhibitor (MNK1 IC_50_ 2.4 nM, MNK2 IC_50_ 1 nM and DRAK1 [STK17A] IC_50_ 82 nM) [25]. Comparing target efficacy (*i.e.,* p-eIF4E inhibition) in mice, ETC-206 is significantly more potent than BAY 1143269. The latter led to only 46% inhibition of p-eIF4E in A549 tumors at approximately at T_max_ at 200 mg/kg at steady state [24], compared to 70% inhibition with 12.5 mg for ETC-206 after a single dose (Fig 3C).

To draw a conclusion on the doses of ETC-206 required for PD inhibition in different murine tissues more weight was given to tumor and skin effects rather than PBMC effects, as the processing time was much faster and hence less variability was observed. The mouse PBMC extraction protocol is lengthy and involves several steps without added phosphatase inhibitor and has previously been demonstrated to show relatively higher variability compared to other murine tissues, while providing much better results in humans [26].

Target efficacy of eFT508 was tested in a B-cell lymphoma xenograft model, where a 1 mg/kg dose gave a 70% inhibition of p-eIF4E up to 8 h, but no inhibition was seen at 16 or 24 h post-dose (at a plasma C_max_ of ∼200 nM), achieving comparable inhibition to ETC-206 at a dose of 12.5 mg/kg. In human patients BAY 1143269 at a dose of 10 mg gave a geometric mean plasma C_max_ of 12.7 µg/L with an AUC_0-24h_ of 169 µg·h/L after a single dose, with dose escalation going up to 50 mg, yielding dose-proportional exposures [27]. ETC-206 at the same 10 mg dose (fasted) in human had a mean plasma C_max_ of 927 µg/L and an AUC_0-t_ of 21,909 µg·h/L [21], demonstrating superior PK properties translated from mice to humans. The development of the BAY 1143269 has since been stopped.

For eFT508, the recommended human dose of 450 mg achieved an exposure of AUC_0-24h_ 7,800 µg·h/L and a C_max_ of 540 µg/L after a single dose [19] with t_1/2_ of 12 h. In the clinic, eFT508 was initially dosed as 450 mg of oral solution once daily, with dosing later changed to a twice daily 200 mg capsule formulation that has improved the AUC at steady state (increase from 5.4 to 10.9 h·µg/mL). At steady state (on Day 15), eFT508 reported a mean 74% inhibition of p-eIF4E in PBMCs for up to 10 h but all other doses tested from 50 mg to 600 mg only lead to mean inhibition of 50-55% [19].

ETC-206 did not show p-eIF4E inhibition in human tissue within the first 6 h after a single dose and it was not expected, based on the C_max_ of 2,000 ng/mL as this did not reach the plasma concentration of 3,500 ng/mL that inhibited p-eIF4E in mouse tissues by ∼70%. However, it is feasible that the long t_1/2_ of ETC-206 and therefore the high AUC_0-t_ of ∼ 60,000 µg·h/L (Fig 6D) contributed to the delayed onset of the p-eIF4E inhibition observed in PBMCs at a dose of 20 mg and to a lower extent at 10 mg. Based on work of Abronzo *et al.* and Eckerdt *et al.* it is also possible that there is a MNK-eIF4E feedback mechanism with contributions by PI3K and mTOR [28, 29] that could cause a delayed PD effect. However, we have no direct experimental evidence for that other than observing rapid initial p-eIF4E inhibition, followed by less inhibition at around 6 h and then more inhibition again between 8-12 h in mice (Fig 2), and seeing delayed inhibition (at 24-30 h) in human samples after a single dose (Fig 4, Fig 5). ETC-206 now enters a clinical trial in cancer patients and it will be best to test for the expected early p-eIF4E inhibition in PBMCs at steady state to provide confirmatory clinical evidence of PD-inhibition. For the clinical development of ETC-206 it was essential to have the p-eIF4E assay sufficiently validated in preclinical species which allowed us to demonstrate that there is trend for delayed p-eIF4E inhibition in human PBMCs.

Here, we have shown that ETC-206 is a selective MNK inhibitor that preclinically leads to dose-dependent PD inhibition in tumors as well as surrogate tissues. In human HVs ETC-206 showed excellent PK properties with a t_1/2_ in humans amenable for every other day dosing and a trend of PD inhibition, which needs to be confirmed in the ongoing Ph2 trial.

## Acknowledgements

This study was funded by the National Medical Research Council, the National Research Foundation of Singapore, and the Biomedical Research Council, A*STAR and scientifically supported by Alex Matter. Shu Jing Lim is acknowledged for her technical support.

## Supporting information

**S1 Table:**
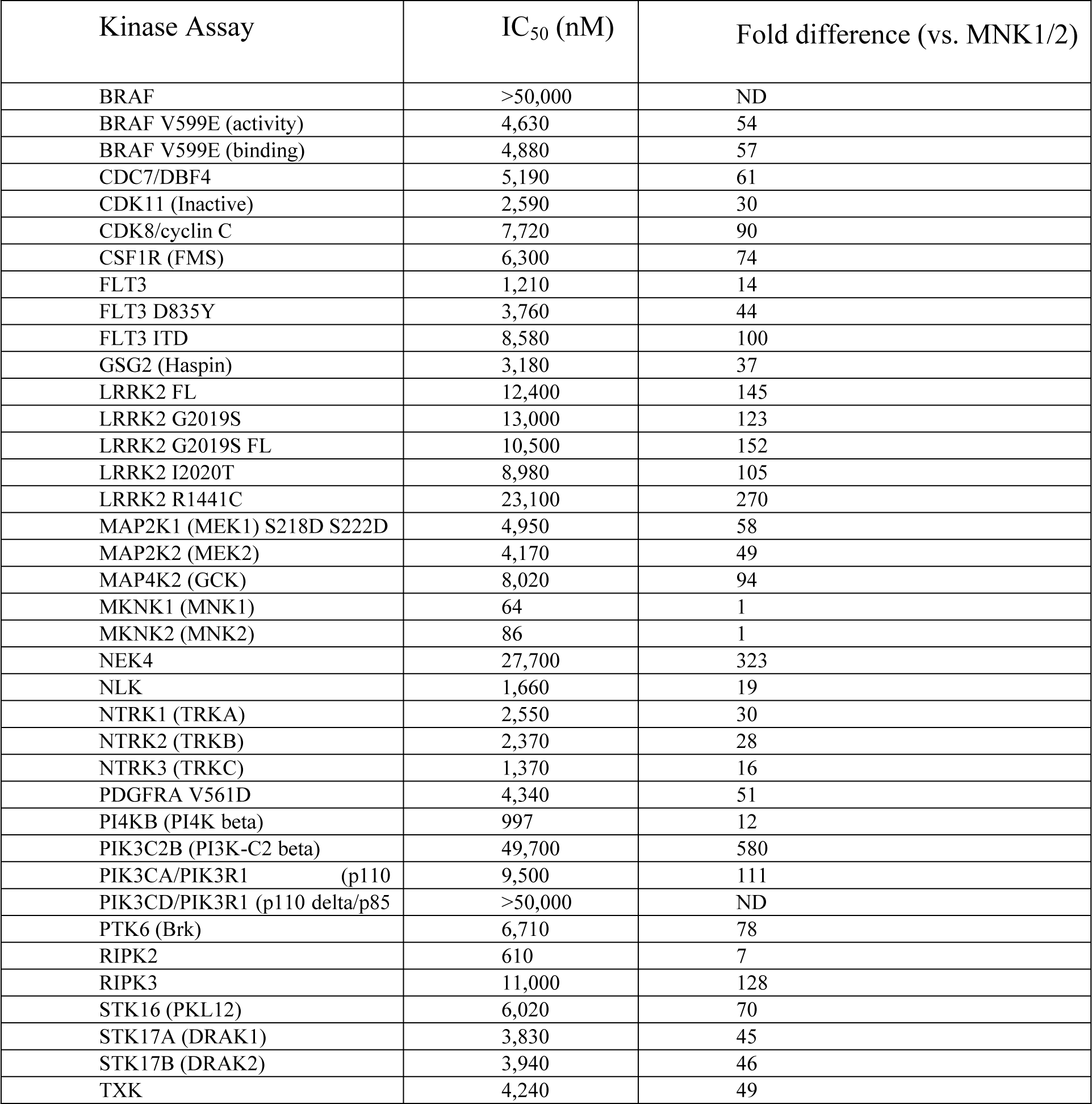
Summary of the IC_50_ values for ETC-206 determined for the 38 primary SelectScreen® kinase hits.

| Kinase Assay | IC<sub>50</sub> (nM) | Fold difference (vs. MNK1/2) |
| --- | --- | --- |
| BRAF | >50,000 | ND |
| BRAF V599E (activity) | 4,630 | 54 |
| BRAF V599E (binding) | 4,880 | 57 |
| CDC7/DBF4 | 5,190 | 61 |
| CDK11 (Inactive) | 2,590 | 30 |
| CDK8/cyclin C | 7,720 | 90 |
| CSF1R (FMS) | 6,300 | 74 |
| FLT3 | 1,210 | 14 |
| FLT3 D835Y | 3,760 | 44 |
| FLT3 ITD | 8,580 | 100 |
| GSG2 (Haspin) | 3,180 | 37 |
| LRRK2 FL | 12,400 | 145 |
| LRRK2 G2019S | 13,000 | 123 |
| LRRK2 G2019S FL | 10,500 | 152 |
| LRRK2 I2020T | 8,980 | 105 |
| LRRK2 R1441C | 23,100 | 270 |
| MAP2K1 (MEK1) S218D S222D | 4,950 | 58 |
| MAP2K2 (MEK2) | 4,170 | 49 |
| MAP4K2 (GCK) | 8,020 | 94 |
| MKNK1 (MNK1) | 64 | 1 |
| MKNK2 (MNK2) | 86 | 1 |
| NEK4 | 27,700 | 323 |
| NLK | 1,660 | 19 |
| NTRK1 (TRKA) | 2,550 | 30 |
| NTRK2 (TRKB) | 2,370 | 28 |
| NTRK3 (TRKC) | 1,370 | 16 |
| PDGFRA V561D | 4,340 | 51 |
| PI4KB (PI4K beta) | 997 | 12 |
| PIK3C2B (PI3K-C2 beta) | 49,700 | 580 |
| PIK3CA/PIK3R1 (p110) | 9,500 | 111 |
| PIK3CD/PIK3R1 (p110 delta/p85) | >50,000 | ND |
| PTK6 (Brk) | 6,710 | 78 |
| RIPK2 | 610 | 7 |
| RIPK3 | 11,000 | 128 |
| STK16 (PKL12) | 6,020 | 70 |
| STK17A (DRAK1) | 3,830 | 45 |
| STK17B (DRAK2) | 3,940 | 46 |
| TXK | 4,240 | 49 |

**S2 Table.** Summary of the IC_50_ values for ETC-206 determined using the CellTiter-Glo^®^ Luminescent Cell Viability Assay 72 h post-treatment for a panel of 71 cell lines, comprising liquid tumor cell lines, lymphoma, myeloma, and non-cancerous human cell lines derived from PBMCs.

| Cell Line | Cell type | IC<sub>50</sub> (μM) |
| --- | --- | --- |
| I9.2 | Acute T-cell leukemia | 6.38 |
| J45.01 | Acute T-cell leukemia | 11.74 |
| P116 | Acute T-cell leukemia | 13.22 |
| J.RT3-T3.5 | Acute T-cell leukemia | 14.84 |
| Jurkat | Acute T-cell leukemia | 17.98 |
| J.gamma.1 | Acute T-cell leukemia | 23.65 |
| D1.1 | Acute T-cell leukemia | >50.00 |
| 8E5 | ALL | 15.32 |
| RS4;11 | ALL | 31.31 |
| KE-37 | ALL | 32.22 |
| Molt-4 | ALL | 39.12 |
| MOLT-3 | ALL | 26.64 |
| CEM-CM3 | ALL (juvenile) | 14.90 |
| CEM/C2 | ALL (juvenile) | 24.04 |
| CCRF-HSB-2 | ALL (juvenile) | 48.67 |
| SUP-B15 | ALL Ph<sup>+</sup> (juvenile) | 38.90 |
| Kasumi-1 | AML | 18.90 |
| Loucy | AML | 33.95 |
| AML-193 | AML | >50.00 |
| MV-4-11 | AML (biphenotypic B-myelomonocytic) | 15.02 |
| EOL-1 | AML (eosinophilic) | 21.43 |
| TF-1 | AML (erythroid) | 48.74 |
| THP-1 | AML (monocytic) | 48.51 |
| BDCM | AML (monocytic, M5a) | 32.21 |
| CESS | AML (myelomonocytic) | 3.15 |
| HL60 | AML (promyelocytic) | 48.50 |
| Clone 15 HL-60 | AML (promyelocytic) | 49.71 |
| BC-2 | B-cell lymphoma | 35.31 |
| BC-1 | B-cell lymphoma | 23.67 |
| MC116 | B-cell lymphoma (undifferentiated) | >50.00 |
| RPMI 7666 | B-lymphoblast (non cancer) | 29.44 |
| GK-5 | B-lymphoblast (non-cancer) | 3.36 |
| AHH-1 | B-lymphoblast (non-cancer) | 26.85 |
| NAMALWA | Bukitt's lymphoma | 1.60 |
| ST486 | Bukitt's lymphoma | 16.03 |
| EB1 | Bukitt's lymphoma | 16.22 |

**S1 Fig.**
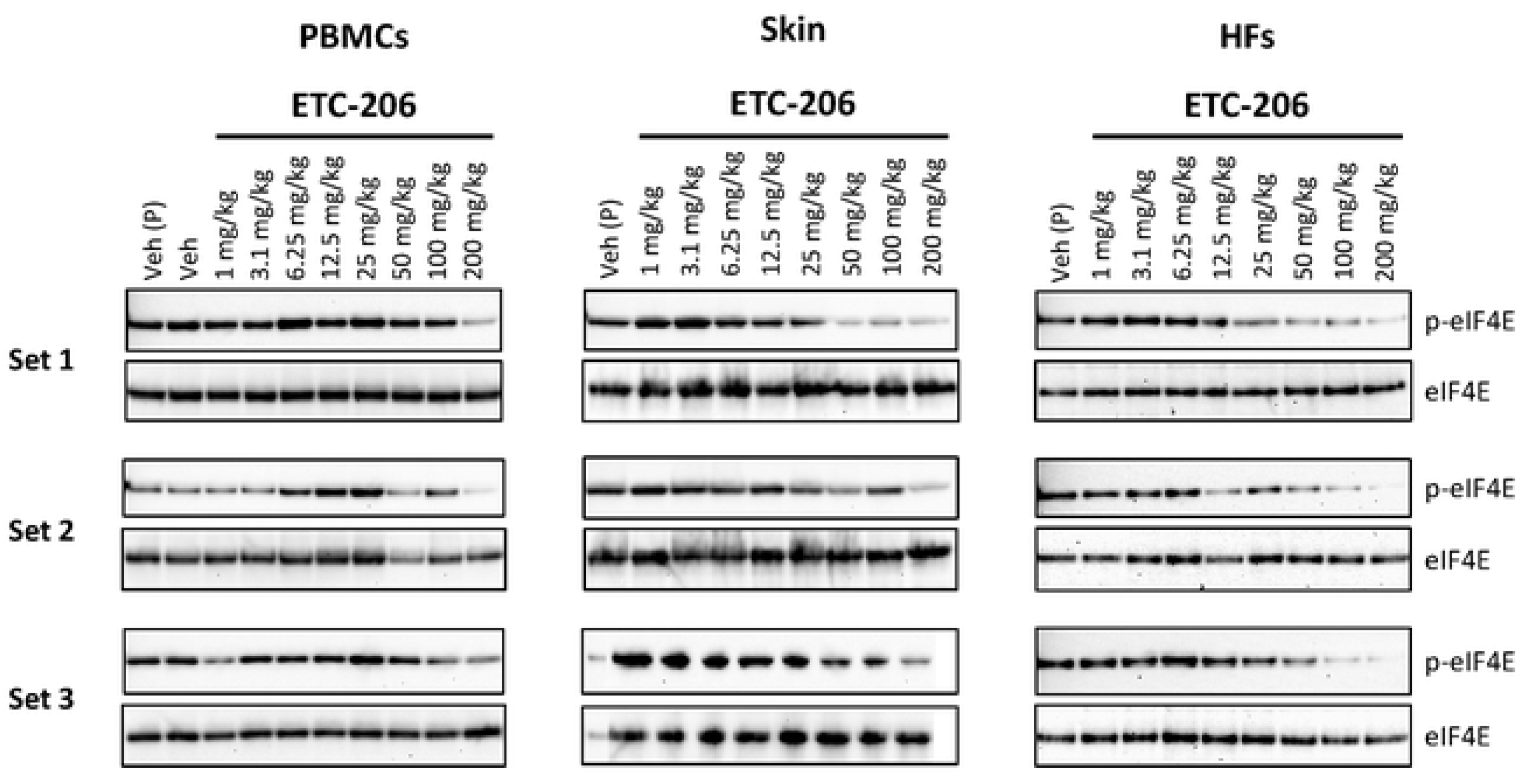
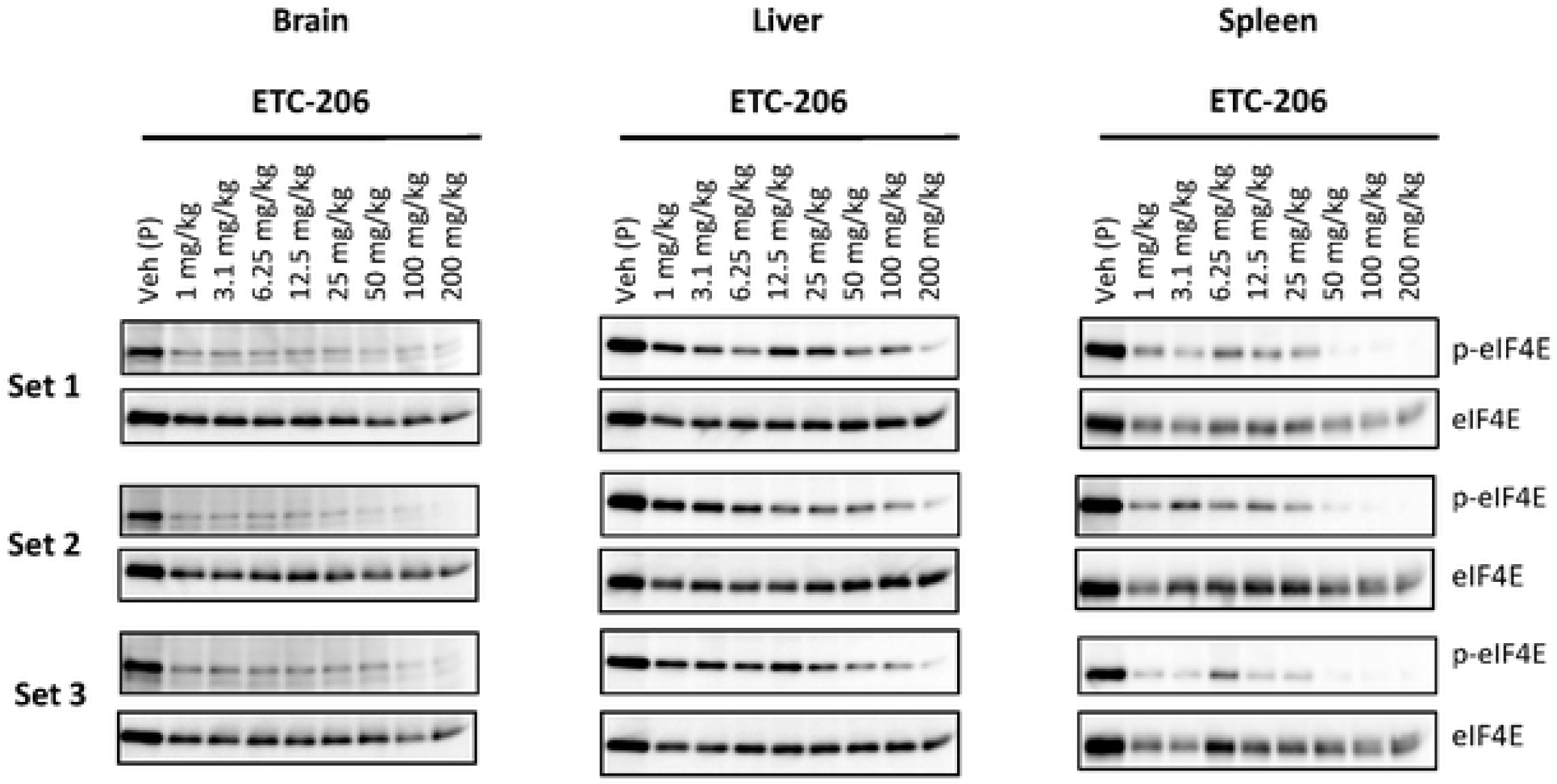
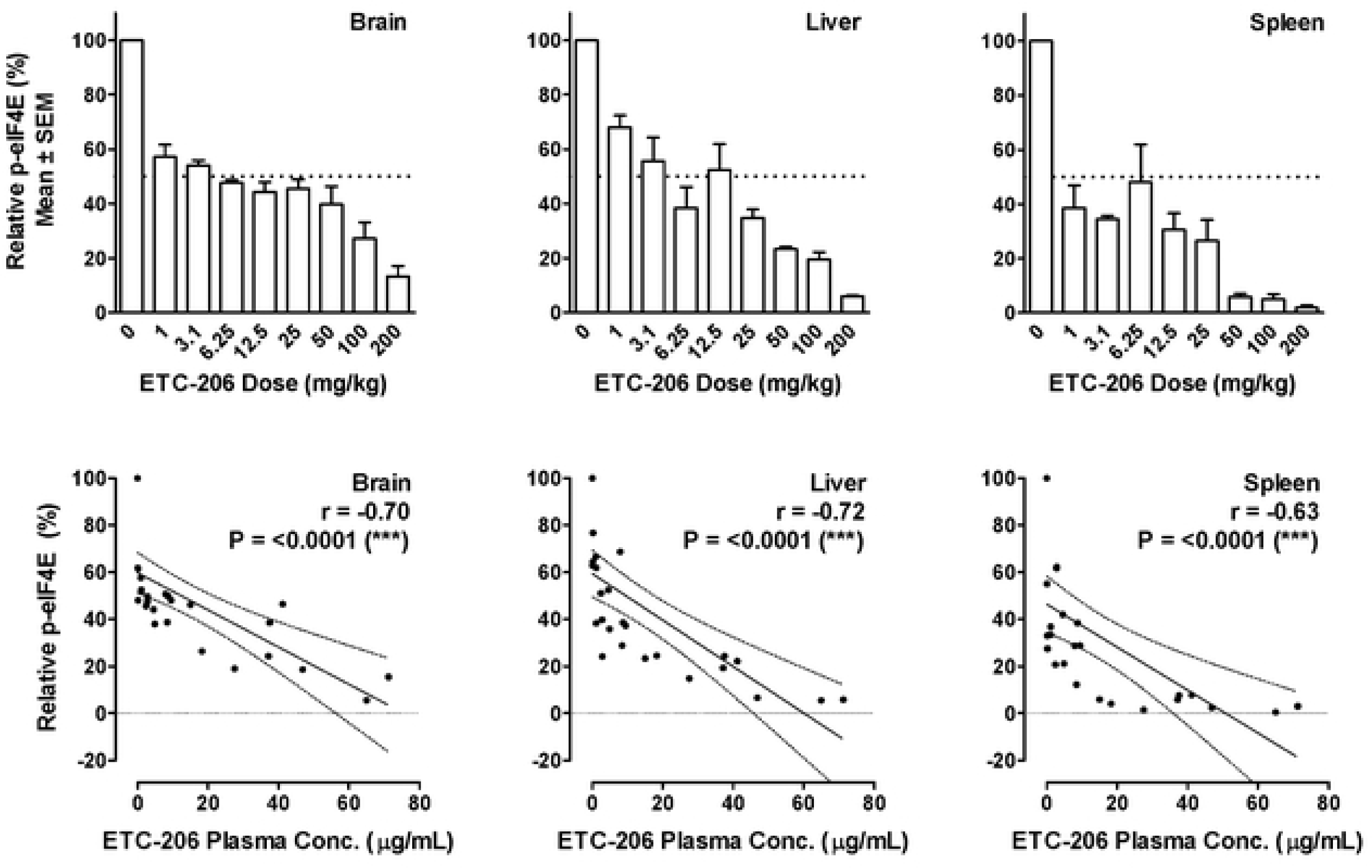
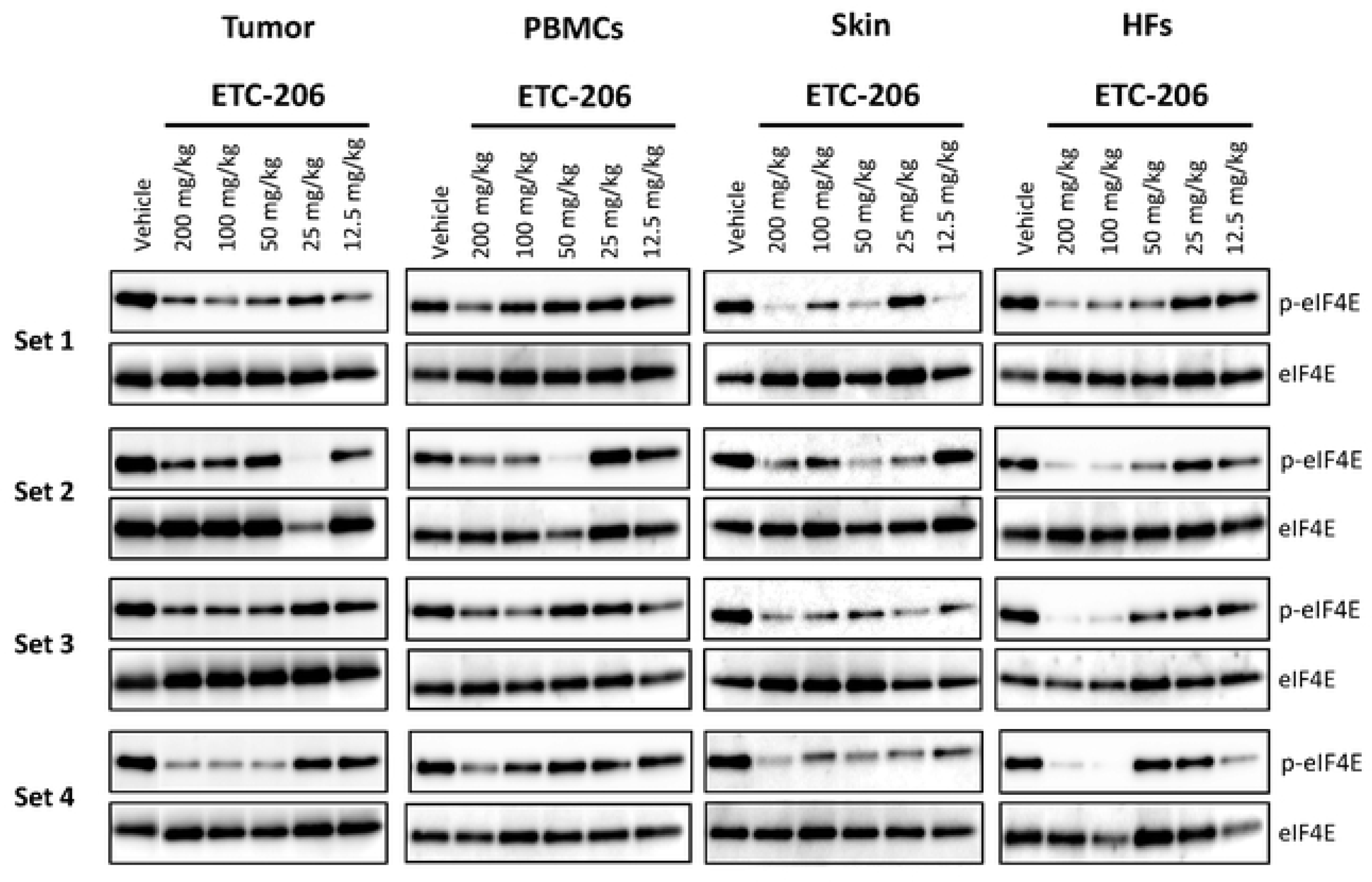
Western blots for p-eIF4E inhibition in different tissues in naïve ICR and tumor-bearing SCID mice after single-dose treatment. (A) Determination of relative p-eIF4E/eIF4E levels in surrogate tissues harvested from female ICR mice treated for 2 h with ETC-206 as indicated. Tissue samples from surrogate tissues were lyzed, separated on Bis-Tris protein gels, and analyzed via Western blot analysis with antibodies as indicated; protein loading was 10 μg of total protein per lane for PBMCs/peripheral blood mononuclear cells, 8 μg of total protein per lane for skin, and 4 μg of total protein per lane for HFs/hair follicles. Veh, vehicle-treated animal, Veh (P), pooled samples from 3 vehicle-treated animals. The corresponding densitometry analysis is shown in Fig 3B. “Set 1”, “Set 2”, etc. represent identically treated sample sets from replicate animals. (B) Brain, liver, and spleen samples (5 μg of total protein was loaded per lane for brain, liver, and spleen) were harvested from the same animals as shown in (A), and analyzed by Western blot as described above. (C) Upper panels: relative p-eIF4E levels were determined from the Western blots shown in (B) and analyzed by densitometry. Lower panels: correlation between ETC-206 plasma concentrations and relative p-eIF4E levels (solid lines: linear correlation, dotted lines: 95% confidence intervals; Pearson’s r and corresponding P values are inserted). (D) Determination of relative p-eIF4E/eIF4E levels in tumor and surrogate tissues harvested from tumor-bearing SCID mice treated with ETC-206 as indicated, and analyzed by Western blot as described above (5 μg of total protein per lane for tumor tissue, 10 μg of total protein per lane for PBMCs/peripheral blood mononuclear cells, 5 μg of total protein per lane for skin, and 3.5 μg total of protein per lane for HFs/hair follicles). The corresponding densitometry analysis is shown in Fig 3C.

**S2 Fig.**
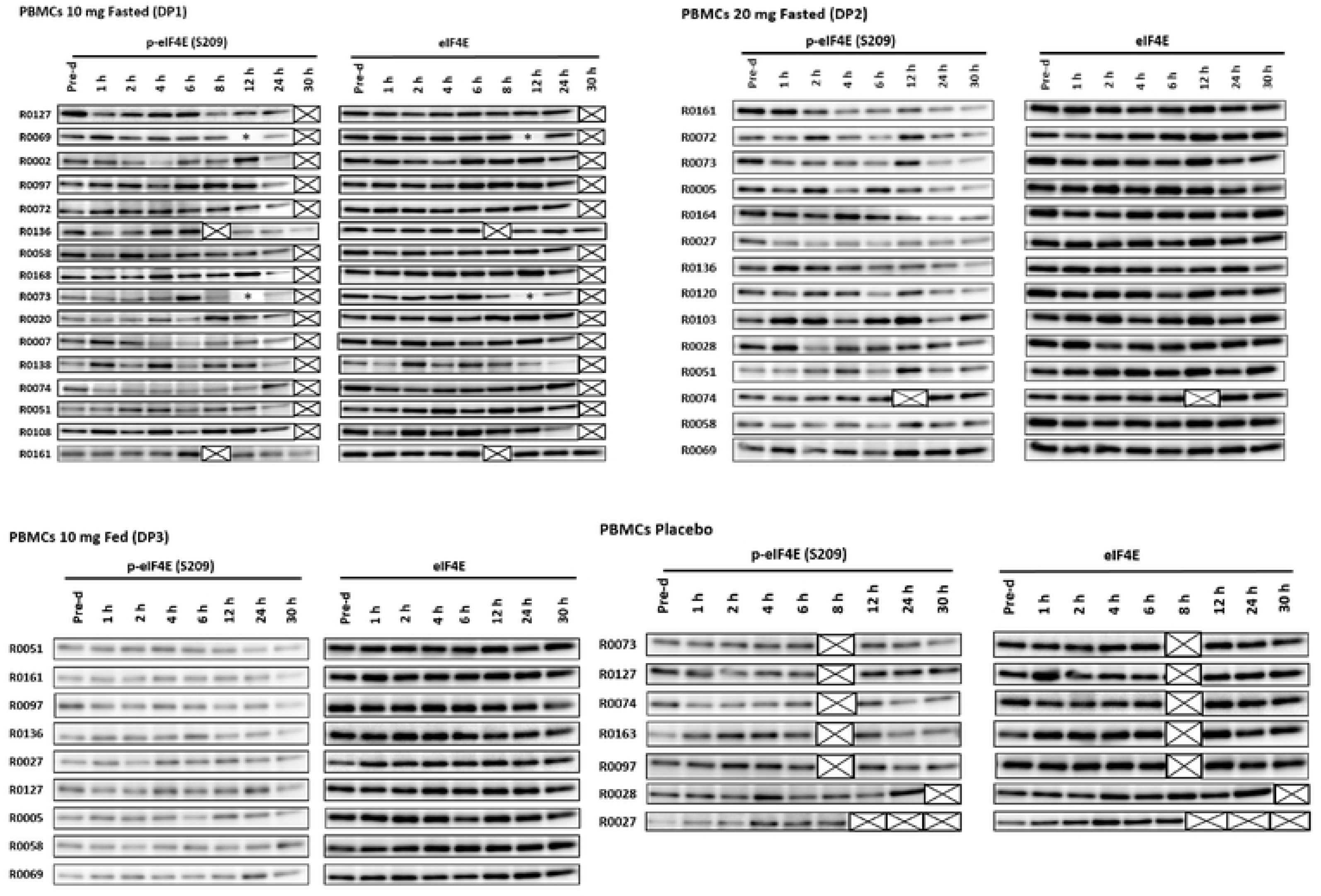
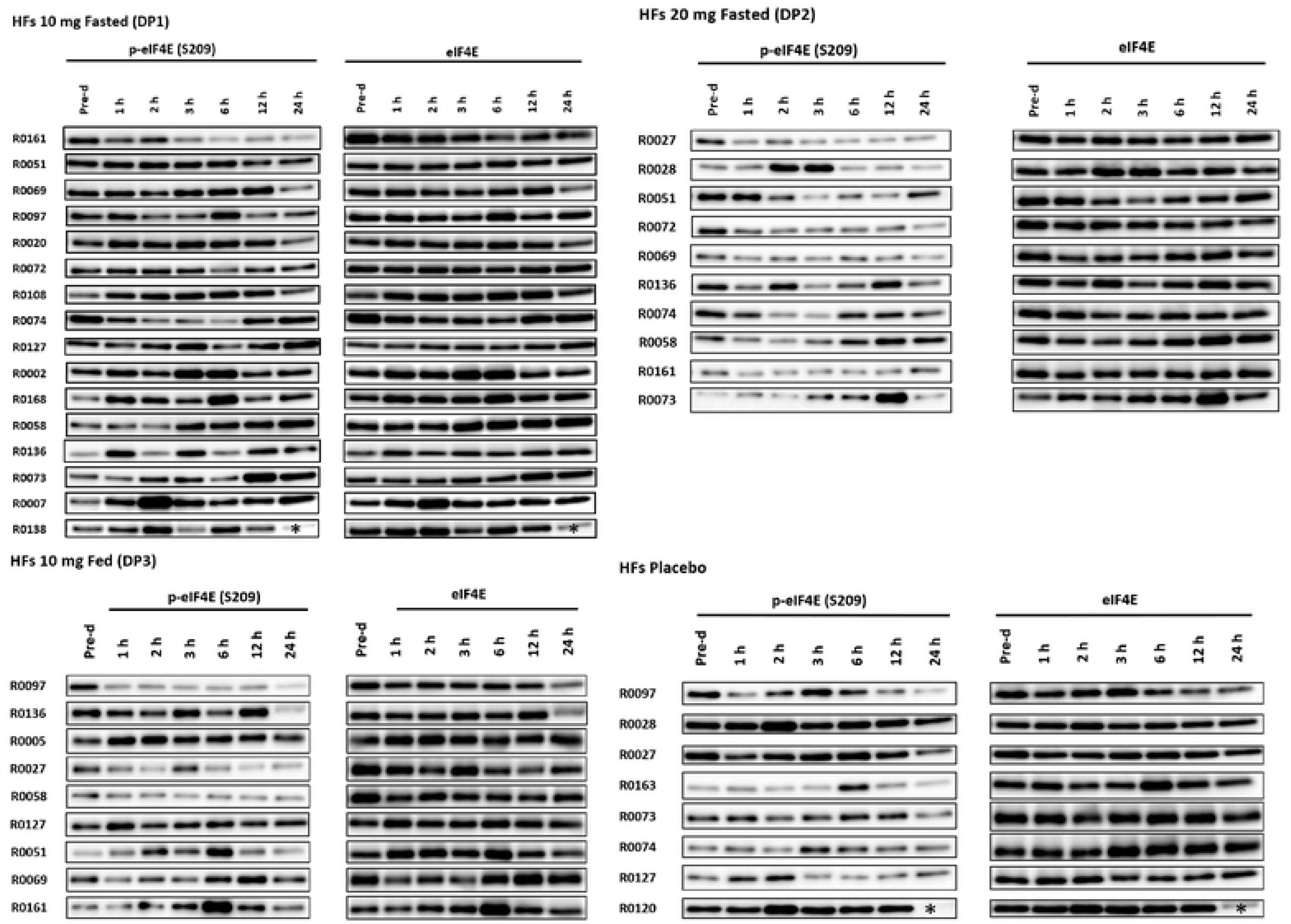
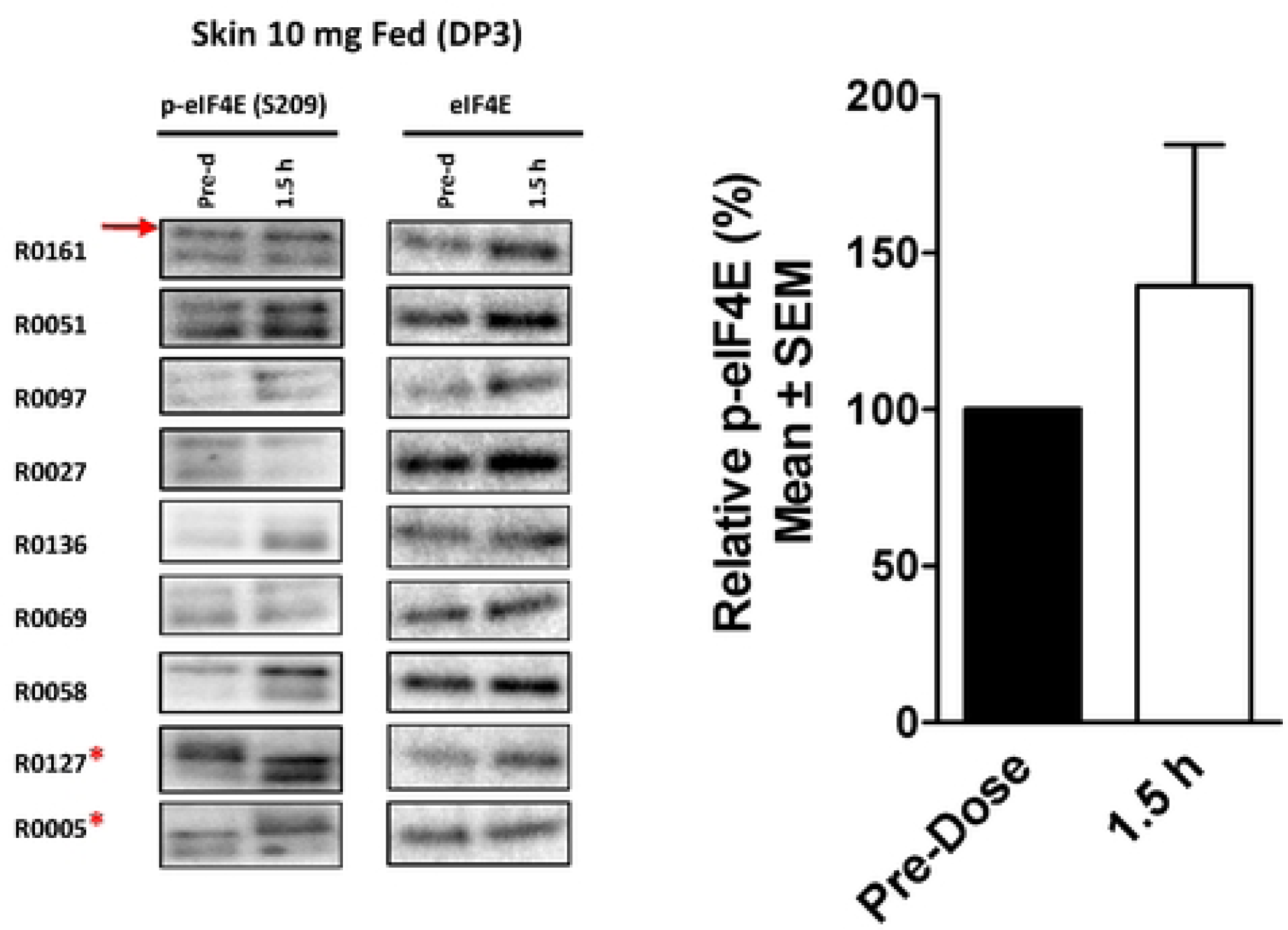
Western blots showing effects of a single-dose of ETC-206 on relative p-eIF4E levels in PBMCs, hair follicles, and skin of human volunteers. (A) Western blots from PBMCs for the data set shown in Fig 4 are shown. Refer to respective figure legend for study/experimental details. Asterisks (*) indicate samples that were partially lost due to cracked vials and values that therefore have not been included, crossed boxes (X) indicate samples that were not collected due to clinical protocol specifications, or outliers. (B) Western blots from PBMCs for the data set shown in Fig 5 are shown. Refer to respective figure legend for study/experimental details. Asterisks (*) indicate samples that were partially lost due to cracked vials and values that therefore have not been included. (C) Effect of a single dose of 10 mg ETC-206 on the relative p-eIF4E levels in skin samples from human healthy volunteers. Subjects received a single dose of 10 mg in the fed state after a high-fat meal. Skin was collected by punch-biopsy from the back from each subject at pre-dose and 1.5 h post-dose (n=9). Western blot results are shown on the left, * indicates samples excluded from densitometry analysis due to incorrect MW; in human skin, a double band is detected of which only the upper band corresponds to the correct molecular weight (red arrow). Mean values of relative p-eIF4E levels at pre- and post-dose are shown on the right (n=7).

## Supplementary Methods

### Mouse blood plasma preparation

Whole blood from mice was obtained by cardiac puncture or retro-orbital bleeding according to local animal welfare regulations, using K_2_EDTA as an anticoagulant. For non-terminal bleedings, ∼0.2 mL blood was collected in 300 µL Microvette® tubes (Cat. #CB300 K2E; Sarstedt, Germany). Blood was centrifuged at 2,000 rpm for 20 min at RT or 4°C. The plasma (supernatant) was transferred into a fresh tube and snap-frozen in liquid nitrogen for further analysis via LCMS/MS-MS.

### Mouse PBMCs

Whole blood (0.7-1 mL) from mice was obtained by cardiac puncture using syringes with 10 μL K_2_EDTA solution and diluted 1:1 with PBS (at RT). The diluted blood was carefully overlaid with 2 mL of Histopaque-1083® (Cat. #10831-100 mL; Sigma-Aldrich) in a 15 mL polystyrene Falcon tube and centrifuged at 400 g at RT for 30 min in a swing-out rotor. The PBMCs were aspirated, washed by addition of 800 µL of cold PBS, and centrifuged at 5,000 rpm in a micro-centrifuge for 1 min at 4°C. The supernatant was removed and the pellet resuspended in 20 - 50 μL of complete lysis buffer. Protein lysates were snap-frozen in liquid nitrogen.

### Mouse Hair follicles

Hair follicles were obtained from alcohol-swabbed skin of mice, preferentially from the mouse whiskers. Individual whiskers were plucked (minimum of 40 whiskers) using sterile forceps, ensuring that the hair root was attached. Plucked whiskers were trimmed at the shafts to ∼1 cm length, if required. A minimum of 40 HFs per time point/dose were collected in a 2 mL homogenization tube (Cat. #TM-625S; Tomy), containing 2 x 2.0 mm Zirconia beads (Cat. #ZB-20; Tomy) and 100 μL of complete lysis buffer. Hair was collected at the bottom of the tube by brief microcentrifugation in a Sorvall Legend Micro 21R (Cat. #75002445; Thermo Fisher Scientific) at 14,000 rpm for 5 min at 4°C.

### Mouse bone marrow

Bone marrow samples were extracted from the left femur of each mouse. The femur of euthanized mice was isolated using macro-dissection, bone marrow was flushed from cleaned femurs using a syringe with an attached 29G needle filled with 2 mL of PBS at room temperature, and the eluate was collected in a sterile 15 mL centrifuge tube. The collected bone marrow cell suspension was diluted with normal saline solution to a final volume of 10 mL. The cell suspension was then centrifuged at 250 g for 8 min. The supernatant was removed and the pellet was resuspended in 200 μl of PBS. In an optional procedure, red blood cells were lyzed using red blood cell (RBC) lysis buffer. To prepare the 10x stock RBC lysis buffer 8.02 g NH_4_Cl, 0.84 g NaHCO_3_, and 0.37 g EDTA-Na_2_ were dissolved in 100 mL MilliQ-water. Bone marrow cell suspension were incubated in 1x RBC lysis buffer for 10 min on ice, followed by centrifugation at 500 g for 5 min at 4°C. The supernatant was removed and the pellet was washed with 500 μL PBS (4°C). The wash was repeated a second time and the pellet lyzed in 100 μL complete lysis buffer. Bone marrow cell suspension where red blood cells were not lyzed were kept on ice until all samples were collected, followed by a centrifugation at 500 g for 5 min at 4°C, one wash with 500 μL cold PBS, and resuspension in 100 μL complete lysis buffer.

### Mouse platelets

For the preparation of platelets, whole blood was obtained via cardiac puncture and collected using sodium citrate as an anticoagulant. Samples were centrifuged at 100 g for 10 min at RT without brake in a swing-out rotor. The resulting platelet-rich plasma was transferred to a separate tube. PGE_1_ (Cat. #P5515; Sigma-Aldrich) was added to each sample to a final concentration of 1 μM, and samples were incubated for 5 min at RT. Sodium citrate (400 μL of a 3.8% [w/v] solution) was added for a final volume of approximately 1 mL. The preparation was centrifuged at 400 g for 10 min (without brake) at 4°C. The supernatant was discarded and the platelets were re-suspended in 20 μL complete lysis buffer.

### Mouse skin

A skin sample (approximately 5 x 5 mm area) was collected from the ventral area of the animals after the fur of the mice had been cleaned using 70% ethanol and had been shaved with a hair clipper. The skin was cleaned again with 70% ethanol, followed by a rinse with neutral saline solution for 3 times before collection. Samples were obtained using sharp scissors and collected in a 2 mL homogenizing tube (Cat. #TM-625S; Tomy) containing one 5.5 mm stainless steel bead each (Cat. #SUB-55; Tomy). Approximately 200 µL of complete lysis buffer was added to each tube prior to snap-freezing in liquid nitrogen.

### Human peripheral blood mononuclear cells (PBMCs)

To prepare PBMCs from human whole blood 8 mL venous blood from healthy human volunteers was drawn into a BD Vacutainer CPT™-tube (Cat. #362761; Becton, Dickinson and Company; Franklin Lakes, NJ) containing sodium citrate, inverted 8 times, and centrifuged for 30 min at room temperature (RT) in a swing-out bucket at 1800 g within 2 h of collection. PBMCs were aspirated from the cell layer above the polyester gel using a disposable pipette, and transferred into a 15 mL conical polypropylene screw cap tube containing 10 mL of PBS. PBMCs were washed 2x using 10 mL of PBS each, followed by a 15 min centrifugation at 300 g and RT. PBMCs were transferred into a 1.5 mL microcentrifuge tube in 100-200 µL of PBS after the first wash, snap-frozen in liquid nitrogen and stored at -80°C. For PD analysis in the EDDC laboratory PBMCs were quickly thawed by addition of 200 µL PBS, pre-warmed to 37°C, followed by centrifugation for 1 min at 4°C and 7,200 rpm in a microcentrifuge. After removal of the PBS, the pellet was lyzed in 25-100 µL cold complete lysis buffer, depending on the size of the pellet. Complete lysis buffer was prepared just prior to use.

Sample collections (blood draws) for PBMC isolation in the 10 mg fasted cohort (Dosing period 1[DP1]) were performed pre-dose and at the following post-dose time points: 1 h, 2 h, 4 h, 6 h, 8 h, 12 h and 24 h in the original protocol version. However, all subjects that have received the 10 mg dose in fasted state at a later time point (i.e. replacement subjects that had completed DP1 at a later stage, after DP2 and DP3, according to Protocol V5.0 dated 18 January 2017) had PBMCs collected at the following, modified time points: pre-dose, 1 h, 2 h, 4 h, 6 h, 12 h, 24 h, and 30 h post-dose. For PBMCs prepared from the 20 mg cohort (DP2) and the 10 mg fed cohort (DP3) the collection time points were the same as for the replacement subjects at 10 mg cohort: pre-dose, 1 h, 2 h, 4 h, 6 h, 12 h, 24 h, and 30 h post-dose (± 6 min).

### Human hair follicles (HFs)

Hair follicles were obtained from alcohol-swabbed skin of healthy volunteers, preferentially from eye brows or beard. Individual hairs were plucked using sterile forceps, ensuring that the hair root was attached. Plucked hairs were trimmed at the shafts to ∼1 cm length, if required. A minimum of 40 HFs per time point were collected in a 2 mL homogenization tube (Cat. #TM-625S; Tomy), containing 2 x 2.0 mm zirconia beads (Cat. #ZB-20; Tomy) and 100 μL of complete lysis buffer. HFs were collected at the bottom of the tube by brief centrifugation in a Sorvall Legend Micro 21R centrifuge (Cat. #75002445; Thermo Fisher Scientific), at 14,000 rpm, for 5 min at 4°C at the following time points: pre-dose and 1 h, 2 h, 3 h, 6 h, 12 h, and 24 h post-dose (±15 min) in all 3 DPs.

### Human plasma preparation

To obtain plasma of human healthy volunteers in the Ph1 study, 3 mL of venous blood was collected using 3.0 mL Vacutainer® K_2_EDTA tubes (Cat. #367856; Becton Dickinson), and blood was mixed with the anti-coagulant via gentle inversion for 8-10x. Plasma was prepared by centrifugation in a table-top centrifuge with a swing-out rotor set to room temperature (18-25°C) at a speed of 1300 g for 10 min. The supernatant was transferred into two cryotubes, snap-frozen in liquid nitrogen and stored at -80°C until transport. Blood samples for plasma preparation for PK analysis were drawn at the following times: pre-dose, 0.25 h, 0.50 h, 1 h, 1.5 h, 2 h, 3 h, 4 h, 6 h, 8 h, 12 h, 24 h, 36 h, and 48 h (± 6 min). Three additional time points (30 h, 72 h [both ± 6 min], and 144 h [± 30 min]) were added in Protocol V5.0 dated 18 January 2017 and were included for HVs dosed in DP2 and DP3 as well as for two replacement subjects that had their DP1 (10 mg fasted) after completing DP2 and/or DP3.

### Human skin punch biopsies

Skin biopsies were obtained from consented subjects in DP3 only under local anesthesia with 1% lidocaine for injection, applied subcutaneously, using a 23G needle. A skin punch from the back of the HVs was obtained using a 3 mm diameter skin punch tool after disinfection of the area with an alcohol swab. The skin core was placed in a 2 mL homogenizing tube containing 8 x 2.0 mm zirconia beads (Cat. #TM-625S with Cat. #ZB-20; Tomy) using sterile forceps, and was snap-frozen in liquid nitrogen and stored below -65°C until homogenization. Two skin biopsies were collected from each subject, one on the day before dosing (D-1) and another one 1.5 h (± 30 min) post-dose. All of the DP3 subjects were supposed to receive 10 mg ETC-206 after a high-fat meal. However, the subjects were accidentally randomized to active and control and therefore 2/9 subjects had a skin biopsy collected after placebo treatment.

### Extraction and bioanalysis of ETC-206 in mouse samples

Extraction was done at the Biological Resource Centre, Singapore, in 20 µL of the diluted tissue using acetonitrile with addition of imipramine (20 ng/mL) as internal standard [IS], Samples (25 µL of supernatant) were precipitated by adding 225 µL of acetonitrile containing IS, vortexed for 10 min at 2,000 rpm in a high-speed microplate shaker, centrifuged for 30 min at 4°C and 4,000 g in a 96 well plate, and injected into UPLC-MS/MS. Bioanalysis of ETC-206 using mass spectrometry from mouse studies was performed using UPLC–MS/MS (Waters Acquity I-Class UPLC system coupled with a Xevo TQ-S Triple Quadrupole Mass spectrometer (both Waters Corporation, Milford, MA). To separate ETC-206 and the IS the chromatography was performed using an Acquity BEH C18 column (2.1 mm × 75 mm; 1.7 µm particle size) coupled with an UPLC column in-line stainless steel filter kit (0.2 µm filter). A gradient method was used for chromatographic separation with the mobile phase consisting of A (10 mM ammonium acetate in deionized water), and B (acetonitrile containing 0.1% formic acid) with the following composition: 0–0.2 min (95% A), 0.2–1.7 min (25 % A), 1.7–1.8 min (95% A), 1.8–2.0 min (95% A). The mobile phase flow rate was 0.40 mL/min. The total run time was 2 min with 2 µL injection volume at a column temperature set to 40°C. For the MS analysis the UPLC flow was analyzed using the Xevo TQ-S (Waters Corporation). Electro-spray ionization (ESI^+^) was used as ionization source, detection was by multiple reaction monitoring (MRM) with following transitions for ETC-206 and the internal standard: m/z 409.17→324.12 and m/z 281.16→58.11 respectively. The ESI–MS/MS parameters settings were as follows: capillary voltage: 4 kV; cone voltage, 90V; source temperature, 135°C; desolvation temperature: 250°C; gas flow to cone 150 L/h, desolvation gas flow: 850 L/h (nitrogen). The collision gas was argon and was introduced into the collision cell at a flow rate of 0.15 mL/min. Data acquisition was done using Masslynx 4.1 software and processed using TargetLynx™ Application Manager (both Waters Corporation).

